# Risk-related decision making evokes distinct brain activation patterns in reward evaluation regions in substance use naïve versus non-naïve adolescents

**DOI:** 10.1101/2020.12.24.424370

**Authors:** Goldie A. McQuaid, Valerie L. Darcey, Amanda E. Patterson, Emma J. Rose, John W. VanMeter, Diana H. Fishbein

**Affiliations:** Center for Functional and Molecular Imaging, Georgetown University Medical Center, Washington, DC, USA; The Interdisciplinary Program in Neuroscience, Georgetown University, Washington DC, USA; Bennett Pierce Prevention Research Center and The Department of Human Development and Family Studies, College of Health and Human Development, The Pennsylvania State University, University Park, PA, USA; Frank Porter Graham Child Development Institute, University of North Carolina, Chapel Hill, NC, USA

**Keywords:** Adolescence, Anterior Cingulate Cortex, Decision-making, Insula, Reward, Substance use

## Abstract

Identifying brain and behavioral precursors to substance use (SU) may guide interventions that delay initiation in youth at risk for SU disorders (SUD). Heightened reward-sensitivity and risk-taking may confer risk for SUD. In a longitudinal, prospective study, we characterized behavioral and neural profiles associated with reward-sensitivity and risk-taking in substance-naïve adolescents, examining whether they differed as a function of SU initiation at 18- and 36-months follow-up.

Adolescents (*N*=70; 11.1-14.0 years) completed a reward-related decision-making task (Wheel of Fortune (WOF)) while undergoing functional MRI. Measures of reward sensitivity (Behavioral Inhibition System-Behavioral Approach System; BIS-BAS), impulsive decision-making (delay discounting task), and SUD risk (Drug Use Screening Inventory, Revised (DUSI-R)) were collected at baseline. Baseline metrics were compared for youth who did (SI; *n*=27) and did not (SN; *n*=43) initiate SU at follow-up.

While groups displayed similar discounting and risk taking behavior, SI youth showed more variable patterns of activation in left insular cortex during high-risk selections, and left anterior cingulate cortex in response to rewarded outcomes. SI participants scored higher on the DUSI-R and BAS subscales. Results suggest differences in brain regions critical in the development and experience of SUDs may precede SU and serve as a biomarker for SUD risk.

## INTRODUCTION

Adolescence is commonly characterized as a period of increased risk-taking coupled with heightened reward sensitivity^1,2^. Risk-taking during this evolutionarily conserved developmental period may have positive outcomes^3^. For instance, increased exploratory behaviors occurring during adolescence allow for adaptive risk-taking^4–7^, which facilitates the achievement of key developmental milestones in preparation for the transition to adulthood^8^. However, brain changes that condition adaptive risk-taking also render adolescents vulnerable to risk-taking that leads to negative outcomes, including substance use (SU)^9^.

Early SU initiation is associated with a constellation of other negative risk-taking behaviors and related adverse outcomes^10^, including delinquency or criminal activity^11^, risky sexual behavior^12,13^, physical assault^14^, accidental injury^15^ and death^16^. Given the potential for deleterious outcomes, SU among adolescents has been identified as a global health concern^17^. Critically, while earlier SU initiation is associated with greater risk for development of lifetime SU disorders (SUDs)^18–27^, any delay in SU initiation decreases risk for development of SUDs^19,28^. For instance, each year of delayed alcohol initiation is associated with a 5-9% decrease in risk for alcohol use disorder^20^. Identifying factors which facilitate prediction of early initiation, therefore, may be beneficial in targeting prevention efforts to delay SU onset.

Developmental neuroscience models offer potential explanations for increased risk-taking during adolescence that leads to SU initiation. The dual systems^29,30^, triadic^31,32^, and imbalance models^33,34^, generally postulate that subcortical brain regions associated with reward processing (e.g., ventral striatum, amygdalae) develop earlier than neocortical regions (e.g., prefrontal cortex (PFC)) associated with cognitive control. Development of the PFC and its functional networks, which continues throughout adolescence and into early adulthood^35,36^, is associated with improvements in top-down control of behavior^37–39^. Such a protracted course of development may render the PFC vulnerable to the impacts of abused substances during adolescence^40^, with early SU potentially altering neurodevelopmental trajectories^41–44^ and, ultimately, adversely affecting adult neurobiology and behavior^9,45^.

Given the proximal and distal negative outcomes associated with early SU, it is critical for the design of effective interventions to identify risk markers that *precede* initiation. Adolescents who initiate SU early often demonstrate preexisting heightened impulsivity,^46,47^ sensation seeking^48^ and reward sensitivity^49–51^. Such traits are associated with poor self-regulation of emotion^52^, behavioral dyscontrol^51^, and a relative imperviousness to punishment^53^, along with increased susceptibility to deviant peer influences^54^. However, these associations do not explain individual differences in outcomes nor do they identify mechanisms that may underlie those differences that would otherwise provide malleable targets for intervention. Brain metrics may, in contrast, predict risk for psychopathology with greater specificity and sensitivity than behavioral measures alone^55^. Thus, it is important to understand underlying neurobiology associated with increased risk^56^ and identify neuroendophenotypes concerning risk for and protection from SUDs^57,58^.

To address whether such biomarkers are observable prior to SU initiation, we followed SU-naïve early adolescents (*N*=70; aged 11.1-14.0 years) over 36 months, and compared participants who did and did not report initiation of alcohol and/or drugs at follow-up. Our central aim was to characterize behavioral and neural profiles associated with reward-sensitivity/risk-taking in relation to SU initiation. Adolescent participants completed measures probing reward sensitivity and risk aversion (Behavioral Inhibition System/Behavioral Activation System (BIS/BAS) Scales), and tasks to assess risk-taking and impulsivity in the context of rewards (Wheel of Fortune and delay discounting tasks, respectively). We predicted that at baseline, those who would go on to initiate SU at 18- or 36-months follow-up would demonstrate greater hedonic and behavioral responsivity to rewards, overvalue immediate rewards, and make riskier choices in a reward-related decision-making task compared to adolescents who remained SU-naïve throughout the study. Further, we predicted that, prior to SU onset, reward-based decision making would be associated with differences between subsequent SU initiators and non-initiators in brain regions implicated in decision-making under uncertainty (i.e., medial prefrontal cortex^59–62^), and in the modulation of reward processing/sensitivity (i.e., ventral striatum and amygdalae^59,63^).

## METHODS

Participants were recruited as part of the Adolescent Development Study (ADS), a prospective longitudinal investigation of the neurodevelopmental precursors to and consequences of early SU initiation and escalation. Detailed information on ADS study methods and aims is presented elsewhere^64^. Briefly, a total of 135 typically developing, SU-naïve early adolescents were recruited from the Metropolitan Washington D.C. region and followed longitudinally. Demographic, cognitive, behavioral, and imaging assessments were conducted at an initial (“baseline”) visit and during two follow-up visits, approximately 18- and 36-months later. The mean time elapsed between the baseline visit and the planned 18-months visit (Wave 2) was 18.4 (SD=3.6) months; and the mean time between baseline and 36-months follow-up (Wave 3) was 36.7 (SD=4.4) months. Imaging and behavioral data reported here were collected during the initial SU-naïve baseline assessment. Exclusionary criteria for the study included adolescent self-report of alcohol (>1 full drink of alcohol at any time) or, with the exception of nicotine, any SU prior to the initial visit; *in utero* exposure to alcohol or illicit drugs (parent-reported); a diagnosed neurodevelopmental disorder (e.g., Autism Spectrum Disorder); left-handedness; a sibling of a current participant; history of head injury resulting in loss of consciousness >5 minutes; or MRI contraindication. The Georgetown University IRB approved all procedures, and written consent and assent were obtained from the parent and adolescent, respectively.

### Participants

Of the 135 participants enrolled in the study, 70 adolescents aged 11.1–14.0 years (*M*=12.7 years, SD=0.66; female=40 (57%)) were included in the analyses reported here. One enrolled participant was excluded due to neurodevelopmental disorder. Participants were excluded from analyses due to missing or incomplete imaging data (*n*=15) and/or excessive head motion during imaging (*n*=24). Additionally, since a primary aim was to examine neural activation during risk-taking, participants who did not make any ‘high-reward/risk’ selections during the Wheel of Fortune task (WOF; see below for paradigm description) were excluded from analyses (*n*=4). Groups were defined based on SU status at follow-up, as detailed below. Participants for whom SU status could not be determined due to attrition or survey discrepancies (*n*=21) were also excluded from analyses reported here. (Table S1 provides a detailed summary of exclusions/inclusions.)

### Family/caregiver measures

#### Socioeconomic (*SES*) *index*

An SES Index was calculated by averaging the mean of two standard scores (mean household income bracket before taxes and mean cumulative years of parental education), and re-standardizing these to obtain a z-score distribution with a 0-centered mean and a standard deviation of 1 for the sample analyzed (*N*=70) (method adapted from^65^).

#### Adolescent measures

##### SU initiation status

At baseline and at the Wave 2 and Wave 3 follow-up visits, adolescents completed two self-report surveys to determine SU status: the Tobacco Alcohol and Drug (TAD) survey and the Drug Use Screening Inventory Revised (DUSI-R)^66,67^. The study-specific TAD included the alcohol and drug portion of the Semi-Structured Interview for the Genetics of Alcoholism^68^ and asked about the use of substances, including tobacco, alcohol, and illicit drugs (i.e., marijuana, cocaine, methamphetamine, ecstasy, opiates, salvia, synthetic marijuana, inhalants, and illegally used prescription drugs), along with an open-ended “any other substances” question. Adolescents also completed the DUSI-R, a survey with demonstrated psychometric validity^69–71^and reliability^72^ for assessing SU and factors associated with risk for SUD later in adolescence. The DUSI-R includes 20 questions concerning use of specific substances (e.g., alcohol, marijuana, prescription painkillers, smoking tobacco, chewing tobacco) or substance classes (e.g., over the counter medications, tranquilizer pills, stimulants).

For the purposes of the analyses reported here, affirmative SU responses on both the TAD *and* the DUSI-R were used in determining SU status. Participants who reported SU on both the TAD and DUSI-R at either Wave 2 or Wave 3 follow-up were categorized as SU initiators (SI). Those who reported no SU on both the TAD and DUSI-R at both follow-up assessments were categorized as SU non-initiators (SN). As detailed above, participants for whom SU status could not be determined were excluded from analyses reported here.

##### DUSI-R absolute problem density (*APD*) *score*

In addition to questions concerning SU, the DUSI-R probes experiences and behaviors known to precede and co-occur with SU. The survey includes eight domains comprised of 159 *yes-no* items that are relevant for early adolescents: SU, behavior, health, social competence, psychiatric symptoms, school performance, family and peer relationships, and recreation^73^. An absolute problem density (APD) score, which reflects overall risk for SU, is calculated by dividing the total number of “yes” questions by the total number of DUSI-R items. Here, group comparisons were conducted for the DUSI-R APD score only.

##### Delay Discounting (*DD*) *task*

Adolescents completed the DD^74^ task outside of the scanner. The task was implemented in E-Prime 2.0. Participants were instructed to choose between receipt of an immediate reward of variable value (≤$10, in increments of $0.50), versus receipt of $10 after a specified temporal delay (e.g., *Would you rather have $2 now, or $10 in 30 days*). Discounting was assessed at six delays: 1, 2, 10, 30, 180, and 365 days. Participants were instructed to make their selections with care, as they would receive a reward (≤$10) based on a random selection of one of their choices^75^.

Values for which the participant demonstrated no preference for immediate versus delayed receipt (i.e., the ‘indifference point’) were normalized to the static delayed reward value ($10)^76^ and plotted against each delay. To adjust for unequal weighting of indifference points at longer delays (a limitation of conventional methods of calculating area under the discounting curve; AUC), while preserving the notion of subjective experience of time via delay scaling (an appeal of conventional AUC metrics), data were log10-transformed (AUClogd^77^). Values ranged from 0 to 1, with smaller AUClogd values representing steeper discounting and thus preference for immediate (smaller, sooner) reward.

##### Behavioral Inhibitory System/Behavioral Activation System (*BIS/BAS*) *Scale*

Adolescent participants completed the BIS/BAS^78^. The BIS/BAS is a 20-item self-report measure answered on a 4-point Likert scale. The BIS is a single, 7-question scale that probes behavioral and emotional responsivity to punishment. Conversely, the BAS is comprised of 3 subscales: Reward Responsiveness, Drive and Fun Seeking. A higher BIS score reflects aversion to and avoidance of potential punishment, while higher BAS subscale scores reflect positive emotionality (Reward Responsiveness) and behavioral approach (Drive and Fun Seeking) in the context of potential rewards.

##### IQ and pubertal development measures

Full-scale IQ (FSIQ) was estimated using the Kaufman Brief Intelligence Test (KBIT), Second Edition^79^. Adolescents completed the Pubertal Development Scale (PDS)^80,81^ as a proxy assessment of physical development via Tanner stage^80^.

##### Wheel of Fortune (*WOF*) *Task*

The WOF task was completed during functional neuroimaging. This well-validated paradigm has been used to probe the neural bases of reward responsivity and risky-decision making under conditions of probabilistic reward versus penalty in both adults^60,61,82,83^ and adolescents^60,83,84^. A modified version of this task was used in this study to probe reinforcing outcomes (i.e., winning or losing) (Figure 1; see Supplement, section S1.1, for further description of the WOF task). Participants were guided through an in-scanner practice (during the structural MRI scan) prior to the actual task to ensure their understanding of how to perform the task. Prior to each run, participants were encouraged to maximize their hypothetical gains and/or exceed their previous total winnings. The task was implemented in E-Prime 2.0, and stimuli were presented via back-projection onto a screen viewed in a mirror mounted to the head coil. A slow event-related design with temporal jitter provided by a variable inter-trial fixation of 2500–10,000 ms based on a Poisson distribution was utilized.

**Figure 1.**
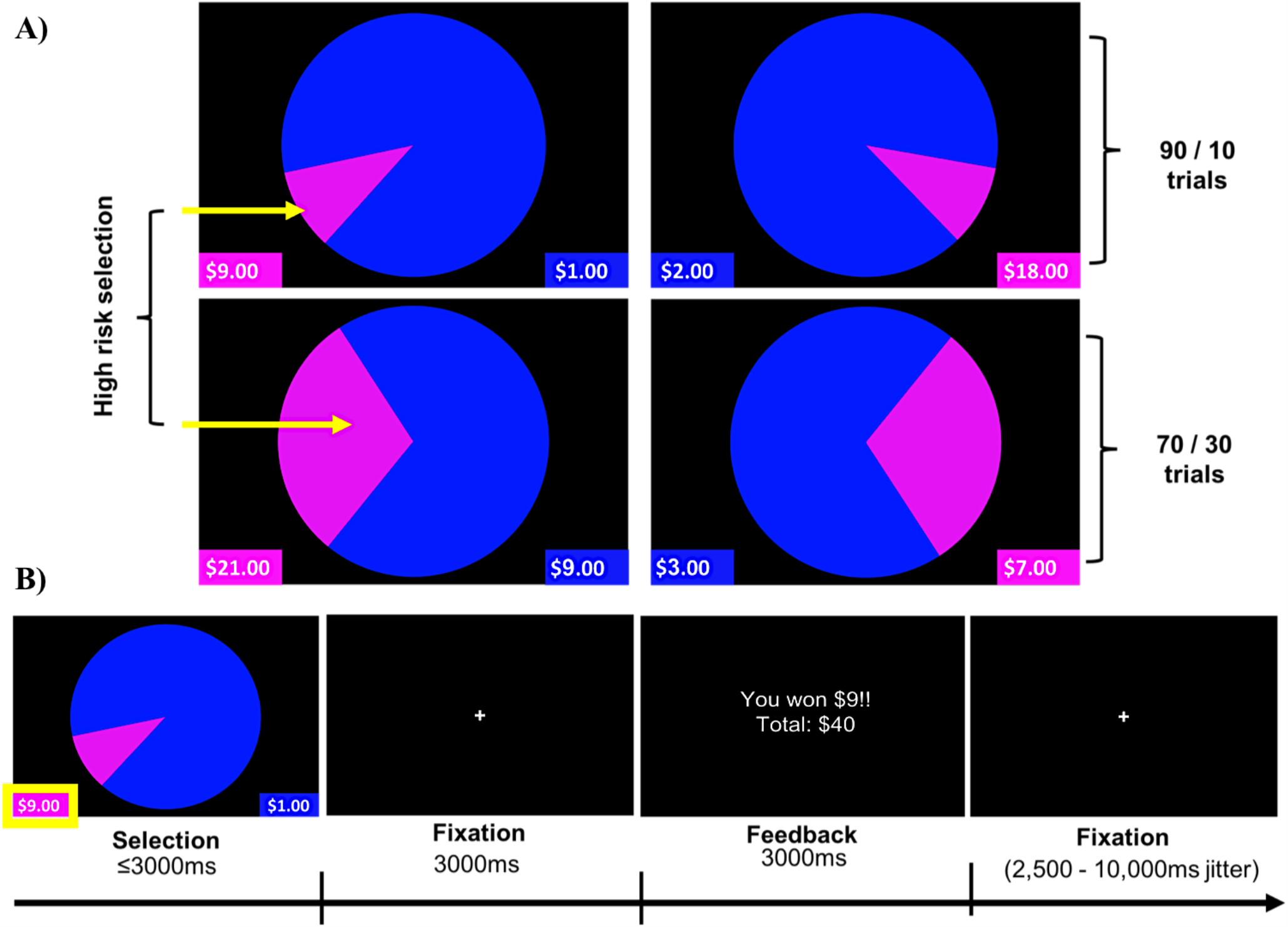
Wheel of fortune (WOF) task. (A) Task stimuli illustrating two trial types (90/10 and 70/30), and highlighting high-reward/risk selections for each of these trial types. (B) Example of a single trial and its timing. Pressing the left button, corresponding to the smaller, magenta portion of the wheel, represents selection of the high-reward/risk option (10% chance of receiving $9), over the low-reward/risk option depicted in blue (90% chance of receiving $1). Analyses reported examined Selection and Feedback phases of the task.

Contrasts of interest for the selection and feedback phases were High-reward/risk>Low-reward/risk and Win>Lose, respectively. Behavioral data analyses considered the percentage of high-reward/risk selections and the average response times (RT) for high-reward/risk and low-reward/risk selections as well as the average RT across all selections.

### MRI Protocol

#### Data acquisition

During the baseline visit, structural and functional images were acquired on a Siemens TIM Trio 3T scanner using a 12-channel head coil. During three runs of the WOF task, functional images were collected using a T2*-weighted gradient-echo planar imaging (EPI) sequence (interleaved slice acquisition, 47 axial slices per volume, TR=2500ms, TE=30ms, TA=2.48ms, slice thickness=3mm, voxel size=3.0 × 3.0 × 3.0mm^3^, FoV=192 × 192mm^2^, flip angle=90^°^).

High-resolution structural images were obtained using a T1-weighted magnetization prepared rapid acquisition gradient echo (MPRAGE) sequence (176 sagittal slices: TR/TE/TI=920/2.52/900 ms, flip angle=9^°^, slice thickness=1.0 mm, FOV=250 × 250 mm^2^, matrix of 256 × 256 for an effective spatial resolution of 0.97 × 0.97 × 1.0 mm^3^).

#### fMRI Data Preprocessing

Image preprocessing and statistical analyses were carried out using SPM8 (http://www.fil.ion.ucl.ac.uk/spm). Preprocessing included correction for interleaved slice timing, realignment of all images to the mean fMRI image to correct for head motion artifacts between images, and coregistration of realigned images to the anatomical MPRAGE. The MPRAGE was segmented and transformed into Montreal Neurological Institute (MNI) standard stereotactic space using non-linear warping. Lastly, these transformation parameters were applied to normalize the functional images into MNI space, and the data were spatially smoothed using a Gaussian kernel of 6 mm^3^ FWHM. A scrubbing algorithm utilizing framewise displacement was implemented to assess participant movement during the fMRI scans^85^. Participants included in analyses demonstrated less than 1mm displacement in fewer than 20% of their total volumes across all three runs of the task.

### Statistical Analyses

#### Imaging data

First-level statistical analyses of imaging data included regressors encoding for trials during which the subject chose either the 10% or 30% probability (High-reward/risk) or the 70% or 90% probability (Low-reward/risk). Regressors of interest also included feedback trials on which subjects won (Win) or lost (Lose). Six translations and rotations modeling participant motion calculated during realignment were included as nuisance regressors.

Contrasts of interest examined whole brain activation for high-reward/risk compared to low-reward/risk trials (High-reward/risk>Low-reward/risk), and winning versus losing outcomes (Win > Lose). Regressors were convolved with the canonical hemodynamic response function. A temporal high-pass filter of 128s was applied to the data to eliminate low-frequency noise (e.g., MRI signal drift). First-level contrasts of interest were used in a second-level analyses for comparisons between SI and SN groups. The initial cluster defining threshold was *p*<.001, with a cluster extent of 10 voxels (voxel size=2.0mm isotropic). Corrections for multiple comparisons were made using a cluster-level FWE threshold of *p*<.05. Macro-anatomical labels reported are based on peak coordinates and were assigned by the Harvard-Oxford Cortical/Subcortical Structural atlases^86–89^, supplemented with labels from *Atlas of the Human Brain, 4*^*th*^ *edition*^90^.

#### Demographics and behavioral data

Statistical analyses were performed using R v.3.5.1. Dependent variables were free from outliers and normality was examined. In the SN group, SES was negatively skewed (Shapiro-Wilk *W*=.88, *p*<.001). BAS fun-seeking scores were non-normally distributed in both groups (SI: *W*=.876, *p*<.05; SN: *W*=.934, *p*<.05); BIS was non-normal for SI (*W*=.897, *p*<.05) and BAS reward responsivity for SN (*W*=.923, *p*<.05). Further, the percent of high-reward/risk selections in the WOF task was positively skewed in both groups (SI: *W*=.757, *p*<.001; SN: *W*=.873, *p*<.001).

Standard transformations for the above dependent variables did not correct distributions; thus, the between-group comparisons were performed using non-parametric Mann-Whitney U test. Alpha was set at *p*=0.05, and Bonferroni correction for multiple comparisons was applied where noted (e.g., BAS adjusted statistical threshold of .05/3=.0167).

With the exception of the group comparisons for DUSI-R APD, all statistical tests were two-tailed. We used one-tailed tests in comparing groups on this measure given a priori evidence of directionality (i.e., DUSI-R APD severity, reflected by higher scores, positively predicts SU^67^).

## RESULTS

### Demographics

SI and SN groups were similar for age, sex, PDS, SES, and race/ethnicity (Table 1). Although mean IQ was lower in SI than SN youth, the difference did not reach statistical significance (*t*(68)=1.92, *p*=.059). Based on reported associations between cognitive ability and measures of decision-making as well as risk-taking^91–95^, we explored whether FSIQ was associated with risk-taking behavior on the WOF task. Given that the percentage of high risk decisions across all runs of the task was positively correlated with IQ (*r*_s_=.28, *p*=.021) (see Figure S1A, Table S2), an effect driven by the percent of high risk decisions in Run 1 (Figure S1B), we treated IQ as a covariate of no interest in imaging analyses presented here. Imaging results for analyses without IQ as a covariate are presented in the supplementary materials (section S2.1, Table S5. Figure S2).

**Table 1.** Participant characteristics at initial assessment

|  | **All**<br>**participants**<br>**N=70** | **SI**<br>**n=27** | **SN**<br>**n=43** | **Test**<br>**statistic** | **p** |
| --- | --- | --- | --- | --- | --- |
| **Age at scan** |  |  |  |  |  |
| Mean(SD) | 12.7(.66) | 12.9(.61) | 12.7(.69) | $t(68)=-1.57$ | .12 |
| Range | 11.1-14.0 | 11.5-14.0 | 11.1-14.0 |  |  |
| **Sex** | | | | $\chi^2=.045(1)$ | .83 |
| Females, n(%) | 40(57%) | 15(56%) | 25(58%) |  |  |
| Males, n(%) | 30(43%) | 12(44%) | 18(42%) |  |  |
| **PDS** |  |  |  |  |  |
| Mean(SD) | 2.3(.67) | 2.4(.64) | 2.2(.68) | $t(68)=-1.42$ | .16 |
| Females | 2.58(.67) | 2.72(.65) | 2.49(.68) | $t(38)=-1.07$ | .29 |
| Males | 1.94 (.47) | 2.1(.46) | 1.83(.47) | $t(28)=-1.55$ | .13 |
| **Race, n(%)** | | | | $\chi^2(3)=3.75$ | .29 |
| African American | 19(27%) | 6(22.2%) | 13(20.2%) |  |  |
| Caucasian | 42(60%) | 15(55.6%) | 27(62.8%) |  |  |
| Hispanic/Latino | 4(6%) | 3(11.1%) | 1(2.3%) |  |  |
| Other | 5(7%) | 3(11.1%) | 2(4.7%) |  |  |
| **FSIQ** | | | | $t(68)=1.92$ | .059 |
| Mean(SD) | 112.1(15.3) | 107.7(14.4) | 114.8(15.4) |  |  |
| **SES Index z-score** |  |  |  |  |  |
| Median(range) | .22(-2.7-1.45) | .22(-2.0-1.45) | .26 (-2.68-1.36) | $U=599$ | .82 |
| Parental education, years, Mean(SD) | 16.6(2.8) | 16.5 (2.8) | 16.6(2.8) |  |  |
| Household income, Median | 100,000-149,000 | 100,000-149,000 | 100,000-149,000 |  |  |

*PDS=pubertal development scale, FSIQ=full-scale IQ, SES=socioeconomic status.*

Among SI adolescents (*n*=27), 12 (44%) and 15 (56%) reported initiation at Wave 2 and Wave 3 follow-up, respectively. While different in age of initiation (Wave 2: *M*=14.79, SD=.41; Wave 3: *M*=15.78, SD=.70; *t*(25)=-4.3, *p*<.001), initiators at both timepoints were similar in demographic characteristics (sex (𝒳^2^(1)=1.08, *p*=.299); race/ethnicity (𝒳^2^(1)=4.72, *p*=.19); age at initial assessment (*t*(25)=.50, *p*=.62); IQ (*t*(25)=-1.69, *p*=.10); pubertal development (*t*(25)=1.66, *p*=.11); SES (*t*(25)=-1.26, *p*=.22)). (See Supplement section, S2.2–S2.4 for SI group SU details).

### Behavioral Results

#### DUSI-R APD, DD, and BIS/BAS

A one-tailed independent samples *t*-test showed adolescents in the SI group had significantly higher scores on the DUSI-R APD compared to the SN group (*t*(67)=-1.89, *p*=.03) (Table 2), suggestive of increased problematic behavior in domains predictive of a future SUD. The SI and SN groups did not differ in discounting behavior (Table 2) indicating similar preferences for immediate rewards. Compared to the SN group, SI adolescents had significantly higher scores on the BAS Drive (*t*(68)=-2.6, *p*=.012) and Fun Seeking (*U*=362.5, *p*=.008) scales, but did not differ for BAS Reward Responsiveness (Table 2).

**Table 2.** DUSI-R APD, DD, and BIS/BAS results

|  | **All**<br>**participants**<br>**$N=70$** | **SI**<br>**$n=27$** | **SN**<br>**$n=43$** | **Test**<br>**statistic** | **$p$** |
| --- | --- | --- | --- | --- | --- |
| **DUSI-R APD** | | $n=26$ | | | |
| Mean (SD)<br>$N=69$ | 16.04(10.28) | 19(11.5) | 14.25(9.1) | $t(67)=-1.9$ | .031* |
| **DD, AUClogd** | | $n=25$ | $n=36$ | | |
| Mean(SD)<br>$N=61$ | .55(.19) | .54(.19) | .55(.19) | $t(59)=.22$ | .83 |
| **BAS Drive** | 9.7 | 10.7 | 9.1 | $t(68)=-2.6$ | .012† |
| Mean(SD) | (2.7) | (2.4) | (2.7) |  |  |
| **BAS Fun-seeking** | 11 | 13 | 11 | $U=362.5$ | .008† |
| Median(Range) | (4–15) | (6–15) | (4–15) |  |  |
| **BAS Reward Responsivity** | | | | $U=516.5$ | .44 |
| Median(Range) | 18(14–20) | 18(14–20) | 18(14–20) |  |  |
| **BIS** | | | | $U=552.5$ | .74 |
| Median(Range) | 20(12–27) | 20(12–24) | 18(15–27) |  |  |

*Note.* Group comparisons for the DUSI-R APD used a one-tailed independent samples *t*-test. Comparisons for AUClogd and the BAS Drive scale used two-tailed independent samples *t*-tests; and those for the BAS Fun Seeking and Reward Responsivity scales and the BIS were conducted using two-tailed Mann-Whitney *U*-tests. \*= $p < .05$ ; †= $p < .0167$ Bonferroni-corrected threshold for BAS subscales. DUSI-R APD=Drug Use Screening Inventory, Revised, Absolute Problem Density; DD, AUC=delay discounting, area under the curve; BAS=Behavioral Activation System; BIS=Behavioral Inhibition System.

#### WOF task behavior

The groups made similar proportions of high-reward/risk selections (*Z*=.537, *p*=.70) (Table 3). A two-way repeated measures ANOVA was used to examine the effect of group (SI vs. SN) and selection type (high-reward/risk vs. low-reward/risk) on RT. A main effect of selection type was found, with both SI and SN groups taking significantly more time to make riskier selections compared to safer ones (*F*(1,68)=65.7, *p*<.0001). There was no significant main effect of group (*F*(1,68)=.0006, *p*=.98), nor was there a significant group × selection type interaction (*F*(1,68)=.007, *p*=.93).

**Table 3.**
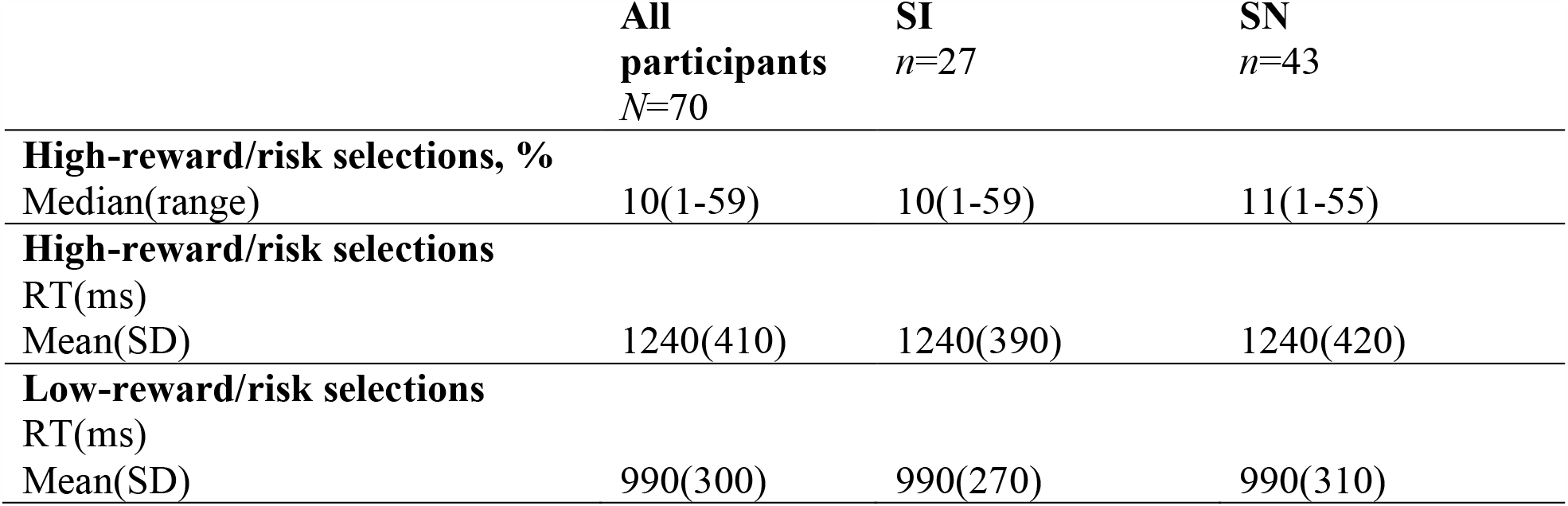
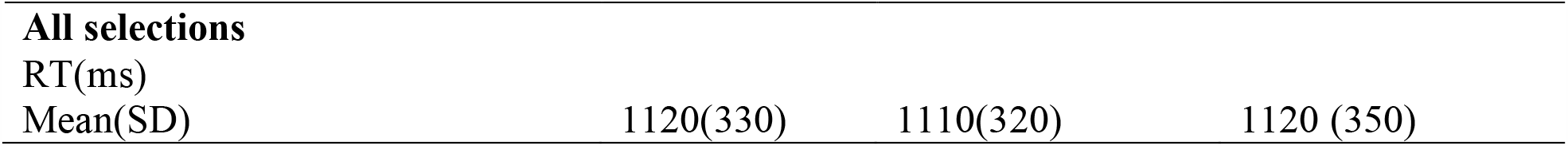
WOF task behavior: descriptive statistics

|  | **All**<br>**participants**<br>***N*=70** | **SI**<br>***n*=27** | **SN**<br>***n*=43** |
| --- | --- | --- | --- |
| **High-reward/risk selections, %** |  |  |  |
| Median(range) | 10(1-59) | 10(1-59) | 11(1-55) |
| **High-reward/risk selections** |  |  |  |
| RT(ms) |  |  |  |
| Mean(SD) | 1240(410) | 1240(390) | 1240(420) |
| **Low-reward/risk selections** |  |  |  |
| RT(ms) |  |  |  |
| Mean(SD) | 990(300) | 990(270) | 990(310) |
| **All selections** |  |  |  |
| RT(ms) |  |  |  |
| Mean(SD) | 1120(330) | 1110(320) | 1120 (350) |

## FMRI results

Compared to SN youth, SI adolescents demonstrated less activation in the left insula when selecting high-reward/risk versus low-reward/risk options. Additionally, when presented with winning versus losing feedback, SI adolescents showed less activation in the left cingulate gyrus, relative to their SN peers (Table 4, Figure 2). There were no regions for which SI youth showed greater activation compared to SN youth for either contrast (see Supplement, Section S21 and Table S5 for presentation of within-group fMRI results). Results surviving corrections for multiple comparisons are presented in Table 4 and Figure 2.

**Table 4.** Summary of SN > SI cluster-level corrected results for two contrasts of interest (IQ as covariate of no interest). Initial cluster defining threshold=p<0.001, k=10 voxels. Reported results survive FWE cluster-correction (p<.05).

| Region | Cluster size | MNI coordinates |  |  | *Z* | *t* | Corrected *p*-value (FWE) |
| --- | --- | --- | --- | --- | --- | --- | --- |
|  |  | x | y | z |  |  |  |
| High-reward/risk > Low-reward/risk |  |  |  |  |  |  |  |
| Left insular cortex | 492 | -34 | 8 | 6 | 3.91 | 4.16 | .001 |
| Win > Lose |  |  |  |  |  |  |  |
| Left anterior cingulate gyrus | 207 | -4 | 20 | 38 | 4.22 | 4.54 | .031 |

**Figure 2.**
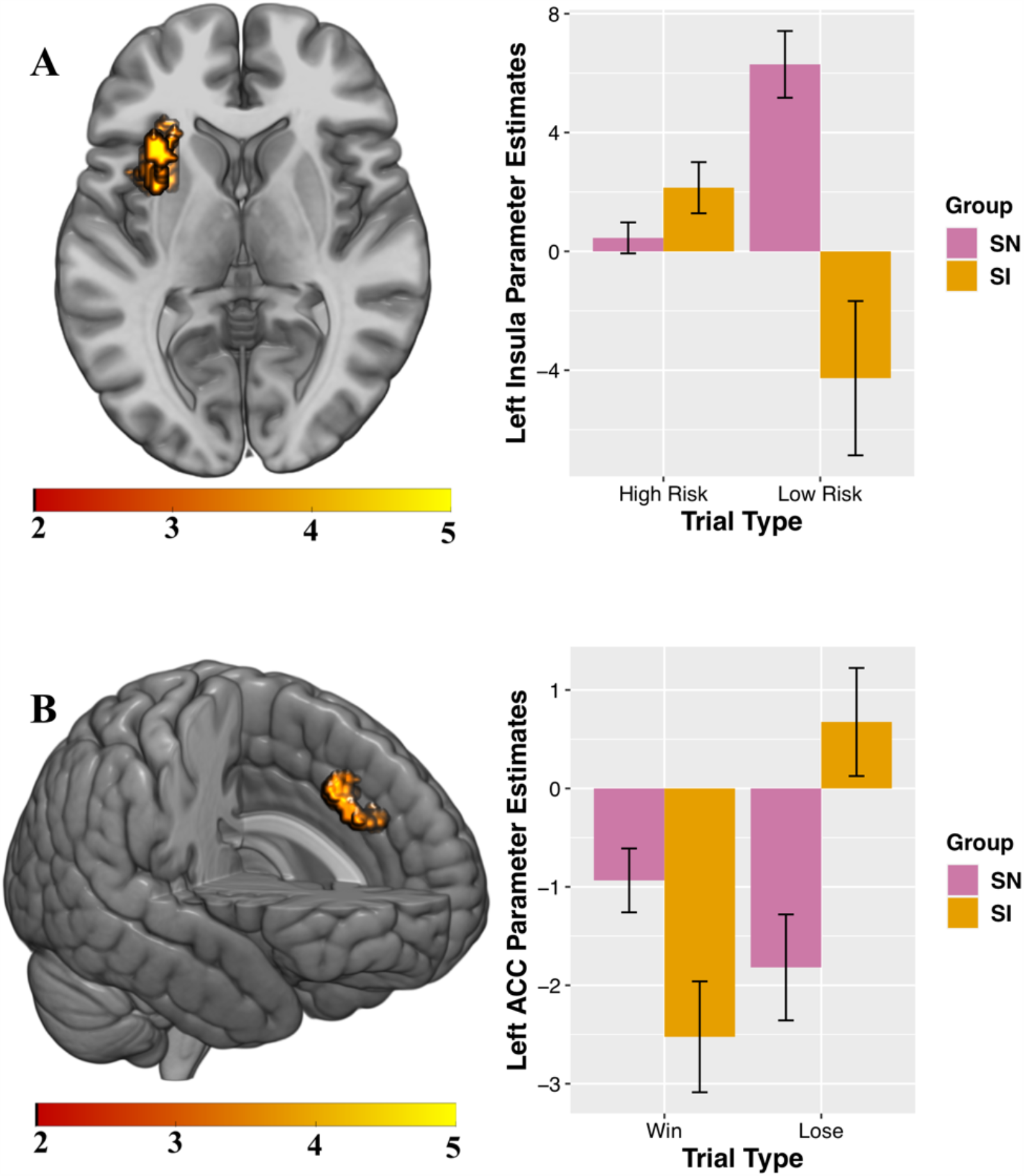
Between-group results for which SN participants demonstrate increased activation relative to SI adolescents. Interaction charts depict mean parameter estimates (error bars represent standard errors) for A) High-reward/risk>Low-reward/risk, left insula; and B) Win>Lose, left anterior cingulate cortex (ACC). FSIQ as covariate of no interest. Initial cluster defining threshold=p<0.001, k=10 voxels. Results survive FWE cluster-correction at p<.05.

### Exploratory analyses: Post-hoc tests of parameter estimates

Visual inspection of the parameter estimates in Figure 2 suggest that the significant between-group results for the contrasts of interest may be driven by differences in how the groups are processing each trial type during selection and feedback. To probe these potential differences, exploratory tests were conducted. All tests were two-tailed. Follow-up independent-samples *t*-tests showed groups significantly differed in activity during low risk (*t*(68)=2.97, *p*=.004) but not high-risk (*t*(68)=-1.25, *p*=.22) trials (Figure 2A). Paired-samples *t*-tests examining within-group processing of high-compared to low-risk trials showed significant differences for SN (*t*(42)=-3.89, *p*=.0004) but not SI youth (*t*(26)=1.95, *p*=.062). Independent-samples *t*-tests showed groups significantly differed for losing (*t*=(68)-2.17, *p*=.034) but not winning (*t*(68)=1.84, *p*=.07) feedback (Figure 2B). Paired-samples *t*-tests revealed SI youth (*t*(26)=-4.35, *p*=.0002) significantly differed for activity when processing winning and losing trial types, while SN do not (*t*(42)=1.59, *p*=.12). Implications of these exploratory analyses as they relate to the between-group findings for High-risk>Low-risk and Win>Lose contrasts are treated in the Discussion.

## DISCUSSION

This study aimed to elucidate neural and behavioral factors that may identify adolescents who are at risk for SU initiation. We examined neural activity during a reward-based decision-making task in SU-naïve early adolescents who either endorsed or denied initiating alcohol and/or drugs during follow-up assessments. Despite similar performance on the risk-taking task, adolescents who went on to report SU initiation showed distinct patterns of activity in left-lateralized insula during risky decisions, and in the left anterior cingulate cortex (ACC) when presented with rewarded outcomes, compared to their non-initiating peers. Furthermore, adolescents who endorsed SU initiation at follow-up reported early increased problematic behavior in domains predictive of a future SUD (DUSI-R APD) as well as greater affective and behavioral responsivity to cues of impending reward (BAS Fun Seeking and Drive) compared to their SN peers. Overall, these findings are consistent with the premise that neural differences in PFC regions may occur *prior* to SU initiation and confer vulnerability to SUDs^96,97^. The findings reported here furthermore lend support to models suggesting divergent neurodevelopmental trajectories may be present prior to SU; as such, due to their malleability, these structures may be targets for early intervention^98^. Critically, the results here underscore the potential utility of neuroimaging in identifying potential precursors of risk for early SU initiation and SUDs.

### WOF task: Behavior and brain

Task-based risk-taking was similar between groups. Compared to low-reward/risk options, selection of high-reward/risk options was accompanied by longer deliberation, an effect consistent with previous studies^61,82,99,100^. Despite equivalent performance, SI and SN groups differed in neural activity underlying *making* risky selections and *processing* rewarded outcomes. The contrast high-reward/risk > low-reward/risk was greater in left insular cortex for SN compared to SI adolescents (Figure 2A). Post-hoc exploratory analyses indicated that during risk-taking SI and SN groups differed in patterns of activation depending on whether they chose a high- or low-risk option, suggesting that the marked difference in responsivity to low-reward/risk trials drives the significant between-group result for the High-reward/risk>Low-reward/risk contrast. Within-group exploratory analyses revealed only SN adolescents showed significantly different activation for high-reward/risk compared to low-reward/risk trials, while activation for the two trial types did not significantly differ for the SI group.

During decision-making, the insular cortex plays a role in refocusing attention based on salience, evaluating risk, inhibiting action, and processing outcomes^101–104^. Attenuated insula activity is associated with increased real-world risk-taking among adolescents^105,106^, and aberrant insula engagement in processing salient stimuli is observed in individuals with addiction^107–109^. Reduced activation of the anterior insula has been found to play an important role in adolescent risky decision making in comparison to adults and is linked to more emotionally driven decisions^110^. Considered within the context of such studies, our results may be indicative of relative immaturity in the SI group in a region that plays an important role in evaluating degree of risk^111^ and as such may have been a contributing factor in the early initiation of substances of abuse in these adolescents.

The contrast Win>Lose was greater for SN compared to SI adolescents in the left ACC. Inspection of parameter plots suggested that while processing outcomes related to gain or loss, SN and SI adolescents demonstrated differing patterns of responses in this region (Figure 2B). Post-hoc exploratory analyses revealed groups significantly differed for activation during losing, but not winning, feedback. Further, only SI youth showed significantly different activation for Lose compared to Win trials. Increased ACC activity has previously been associated with processing gains (relative to no gains) in a gambling task in adolescents^100^. Individuals with established SUDs show impairments in decision-making^112^, altered ACC structure^113^ and differences in brain activity during risk-taking^114^. Specifically, individuals with SUDs display not only greater substance-related cue-induced ACC activity during active use^115^, but also blunted ACC activity during decision-making while abstinent^116^, an effect which predicts craving, length of time to relapse and relapse severity^117^. Importantly, some of these differences may be evident prior to development of AUD/SUD including alterations in ACC neuroanatomy^118^ and suggest increased vulnerability to SUDs (e.g., youth with positive family history of alcoholism demonstrate hyper-activation during risk taking compared to youth with negative family history)^119^.

Both the insula and ACC are implicated in reward-related decision-making^60,82,105,111,120,121^ and the ACC exhibits age-related decreases in activity during risky-decision-making^122^, while adolescents, compared to adults, show increased activation in both ACC and insula during reward processing^123^. Moreover, as hubs in the functional salience network, the anterior insula and ACC^124,125^ are believed to integrate automatic, bottom-up detection of relevant internal and external stimuli with cognitive, top-down processing^126^. The salience network is implicated not only in altered cue-reactivity among individuals with SUD^107,108^, but may play a contributory etiological role in early SU and transition to SUD^127^. While others have established that adults with SUD demonstrate aberrant patterns of insular and cingulate activity during risky decision-making^128^ and that reduced insular activity during risk-related processing is predictive of relapse^129^, our results suggest that variability in insular and ACC activity is present in individuals at risk for SUD even *before* substance initiation.

The exploratory results intriguingly suggest SI youth may be more sensitive to the distinction between wins and losses, though this remains to be empirically tested in planned comparisons correcting for multiple comparisons. It is also possible that steep hypoactivation of the ACC in SI adolescents in the context of rewarding outcomes indicates an increased threshold for rewarding stimuli (consistent with elevated BAS fun-seeking scores SI youth). These group differences may reflect differences in outcome monitoring and processing^130^ and awareness of outcomes^131^, which serve in part to guide behavior^132,133^. A notable consistency between present findings and previous studies is that youth with differential risk for SU demonstrate similar task performance but differences in patterns of brain activation across a variety of tasks^134,135^. Thus, early disruptions in PFC function, including ACC, may contribute to a constellation of impairments including aberrant response inhibition and salience assignment^97^ and ultimately to real-world risky decision making.

### Measures of risk and reward-sensitivity: DUSI-R, BIS/BAS, and DD

Compared to their SN peers, SI adolescents showed elevated DUSI-R absolute problem density (APD) scores. Because the APD score is a multi-dimensional construct, quantifying adolescent difficulties across health, psychosocial and psychiatric domains associated with SUD^66^, elevated APD scores in SI adolescents may reflect increased relative risk for SU. It should be noted, however, that 69% of SI youth would *not be considered high risk* according to the previously established cut-off score of 24^67^; further, similar proportions of adolescents in each group scored ≥24 (SI: 30% (8/27); SN: 19% (8/43); *𝒳*^*2*^=.70(1), *p*=.40). In contrast, the DUSI APD validation study in youth aged 12-14 years predicted SUD (according to DSM-III-R criteria) by age 19 with 73% accuracy^67^. Although early initiation is itself a key risk factor for SUDs, as noted earlier, indices of brain function may be more sensitive and specific indicators of risk than outward behavior and may be apparent *prior to SU onset*. If so, such measures may identify potent targets for effective interventions to prevent SUDs.

Prior to SU, SI adolescents (relative to their SN peers) showed elevated BAS Fun Seeking and Drive scores. Elevated scores on these two BAS scales have been associated with low levels of harm avoidance^78^ and problematic alcohol use, including among adolescents^136–139^. Further, these subscales were positively correlated with adolescent risky choices during a win-only (but not a lose-only) version of a WOF task^99^. Thus, elevated scores on these BAS subscales prior to SU may reflect a propensity towards affective and behavioral responsivity to rewarding stimuli in SI youth, which may bias individuals toward greater real-world situational risk-taking and decreased harm avoidance^101^.

SI and SN youth did not differ on performance on the DD task, suggesting similar preference for immediate rewards under current task parameters. Unlike the BIS/BAS and DUSI-R which assess real-world preference and situationally-based behavior, the laboratory DD task (like WOF) may lack the sensitivity to detect group differences prior to initiation^140^. The absence of differences between SN and SI participants on the DD and WOF task performance, in the context of functional differences during WOF performance, lends further support to the notion that brain indices may be more sensitive to risk prior to the onset of SU, compared to behavioral measures.

## Strengths and Limitations

An important strength of the current study is the stringent inclusion criteria implemented to ensure SU-naïve status of youth at initial assessment, and the requirement of convergent responses on two SU measures (DUSI-R and TAD) to classify youth at follow-up. In contrast, previous studies examining “SU-naïve” youth include those who report “little to no” alcohol use^141^, or who do not report “significant”^59,63,142^ or “heavy” alcohol or drug use^143^. Others rely on urine drug screening at scanning time^84^, which, for many drugs, capture only recent use^144^ and are not reflective of patterns of use over time.

Another strength of the current study is the narrow age range at baseline (11-13-year-olds). While previous studies enrolled participants with a more distributed age range^60,84,142^, we restricted eligibility at enrollment to a much smaller range in an effort to capture information regarding early initiation and minimize potential age-related confounds in neurodevelopment. Finally, the current sample of adolescents were well-characterized using a battery capturing a variety of factors presumed to confer risk for or resilience to early SU, including preference for immediate gratification (DD), affective and behavioral responsivity to rewards and punishment (BIS/BAS), and multidimensional risk for SU problems (DUSI-R).

On the other hand, by analyzing the selection phase of the WOF in a version of the task that consistently coupled high reward with low probability and low reward with high probability we were unable to dissociate between patterns of activation associated with reward versus risk. Although this limitation is not unique to the current study^145^, it is unclear here whether between-group differences in insular cortex were driven by reward sensitivity or risky decision-making. Inclusion of choices with equal probability of high/low reward (i.e., 50/50 wheels), as in the ‘classic’ WOF task, would have permitted testing the relative contributions of reward magnitude independent of perceived risk^61^. It is important to note, however, that estimation of reward value and tendency towards risk outside of the laboratory may not be not entirely separable either; decisions with greater reward potential, whether adaptive (e.g., approaching a classmate to initiate a conversation) or maladaptive (e.g., underage alcohol consumption) are inherently accompanied by risk (e.g., social consequences such as peer rejection; or adverse physiological impacts of alcohol consumption and parental or school punishment for drinking).

The current study identified youth who initiated at different ages (initiation at approximately18- vs. 36-months follow-up), which may also limit the interpretation of our outcomes. Although the two SI subgroups (i.e., earlier vs. later initiators) were similar in demographic, physical and cognitive characteristics, as well as task-based behavior and BIS/BAS scores, those who reported earlier initiation (approximately 18 month follow-up) scored higher on DUSI-APD, indicative of greater risk in domains that precede or co-occur with problematic SU. Due to concerns regarding statistical power, we were unable to compare SI subgroups on brain activation during the WOF task. It is recommended that future studies recruit greater numbers of participants who are likely to be assigned to one of these two SU subgroups, in order to increase the statistical power to prospectively examine group differences in neural activation among individuals who initiate at different stages of adolescence.

Relatedly, the current study is unable to determine pathways to SU escalation, and SUD. SU initiation itself is not necessarily indicative of continued or escalated use or the eventual entrenchment of pathways that might be specific to SUD risk. The elucidation of factors that give rise to such pathways, including early brain biomarkers, may provide a much richer understanding of how brain functioning in SU-naïve adolescents portends subsequent life course outcomes.

## CONCLUSIONS

During adolescence, while exploratory behavior like risk-taking is normative and even adaptive under certain circumstances, the brain changes that drive risk-taking during this developmental period may also facilitate vulnerabilities for maladaptive risk-taking including early SU initiation and escalation. Earlier SU is associated with adverse outcomes and dramatically elevated risk for SUDs, which present a lifelong burden to the individual. Early SU may also alter the neurodevelopment of regions critical to impulse control, further compounding predispositions of youth to risk-taking^33,44,146^. Thus, identification of neuroendophenotypes for early SU initiation is crucial to inform targeted interventions among at-risk youth *prior to* SU entrenchment. At a minimum, delaying SU in these individuals would allow time for cognitive control functions supported by the developing PFC to come on board and may thus confer protection against SUDs in adulthood^127^. The current study corroborates and extends the literature by demonstrating that variability in activity in ACC and insula—key regions known to support reward- and risk-related decision-making—may distinguish SU-naïve youth who initiate SU early from those who remain abstinent.

## Acknowledgements

This work was supported by the National Institutes of Health/National Institute on Alcohol Abuse and Alcoholism [1R01AA019983-01; 3R01AA019983-02S1; 5F31AA023462-02], NIH/NCATS [1KL2RR031974-01, and NIH/NICHD [2P30HD040677-11].

