## Supplemental materials for "Risk-related decision making evokes distinct brain activation patterns in reward evaluation regions in substance use naïve versus non-naïve adolescents"

### S1. METHODS

*Table S1. Participants remaining after each stage of data filtering based on inclusionary/exclusionary criteria*

| | *N* |
|---|---|
| 1. **Enrolled in study** | **135** |
| 2. **Neurodevelopmental disorder** | 1 |
| 3. **Indeterminate use status** | **21** |
| 4. **Excluded due to incomplete or insufficient imaging or task data** | |
| a. No imaging data | 1 |
| b. Technical problems during imaging data acquisition | 10 |
| c. Incomplete imaging data | 4 |
| d. No high-reward/risk selections during task | 4 |
| 5. **Excluded due to excessive head motion** | **24** |
| TOTAL excluded | 65 |
| **Final sample** | **70** |

*S1.1. Wheel of Fortune (WOF) task*

During each trial of a modified WOF task[1], participants were presented with a 'wheel' (a probability pie-chart), divided into two unequal slices, which depicted the odds of winning hypothetical monetary rewards associated with those slices (Figure 1). Across 90 trials (3 x 30 trial runs; approximately 21 minutes in total), the odds of winning/losing were randomly varied between 10% vs. 90% (32-42 trials) and 30% vs. 70% (48-58 trials). Monetary amounts ranged from $1 to $21. During the selection phase of the task, adolescents chose between a relatively small chance (10% or 30%) of winning/losing a large reward ($9, $18 or $7, $21) (i.e., 'high-reward/risk' selection), versus a greater chance (90% or 70%) of winning/losing a smaller reward ($1, $2 or $3, $9) (i.e., 'low-reward/risk' selection).

The present study used a version of the WOF task that was modified to reduce scan time and minimize fatigue for our early adolescent participants. This included excluding trials with equal odds (50/50) of winning either value and the anticipatory phase, during which the participant rated their confidence in their selection outcome. Further, the task was modified so that both ‘wins’ *and* ‘losses’ were possible outcomes.

Participants made a selection by pressing the button on the side corresponding to the color of their choice (Figure 1). Participants were given 3000ms to make a selection, and failure to respond within the allotted time resulted in loss of the highest dollar value offered for that trial. Selections were followed by a 3000 ms delay preceding the feedback phase during which participants viewed a screen informing them of the trial outcome along with a running total of their winnings.

*S1.2. BIS/BAS*

The BIS/BAS is a 20-item self-report measure answered on a 4-point Likert scale (“Very true for me,” “Somewhat true for me,” “Somewhat false for me,” “Very false for me”)[2]. The BIS is a single, 7-question scale that probes behavioral and emotional responsivity to punishment). The BAS, on the other hand, is comprised of 3 subscales: Reward Responsiveness (5 questions, which probe affective responsivity to rewards), Drive (4 questions, which probe persistence in the pursuit of rewards), and Fun Seeking (4 questions, which probe a desire for novel rewards and spontaneity in acting to obtain rewards). A higher BIS score reflects aversion to and avoidance of potential punishment; while higher BAS subscale scores reflect positive emotionality (Reward Responsiveness) and behavioral approach (Drive and Fun Seeking) in the context of potential rewards.

*S1.3 Associations between WOF task behavior and full-scale IQ (FSIQ)*

We examined whether FSIQ was associated with the percent of high-reward/risk decisions and RT for high-reward/risk decisions in WOF task behavior. In the total group ($N$=70) FSIQ was positively correlated with the percent of high risk decisions for all three runs of the task ($r_s$ =.28, $p$=.021) (Figure S1A). Looking more closely at the percent of high-reward/risk decisions by run (Table S2) we found that this correlation is driven by the association of FSIQ with high-reward/risk decisions in the first run of the task ($r_s$=.31, $p$=.0083) (Figure S1B). There were no significant associations between FSIQ and either the second ($r_s$ =.09, $p$=.44) or third run of the task ($r_s$ =.04, $p$=.74). Using the R package WRS2, a robust repeated-measures mixed ANOVA showed only a main effect for Run ($F(2,28)=7.99$, $p$=.002), with no significant effect for group ($F(1, 32)=.17$, $p$=.68) or the run × group interaction ($F(2,28)=.14$, $p$=.87). Mean RT for high-reward/risk decisions was not significantly correlated with FSIQ, neither for all 3 runs nor for any single run (all *p*s >.05).

*Table S2. Percent of High-reward/risk selections on the WOF task by run*

| | SI<br>*n*=27 | SN<br>*n*=43 |
|---|---|---|
| **High-reward/Risk selections, Run 1, %** | | |
| Mean(SD) | 21.8(21.7) | 22.3(20.6) |
| Median(range) | 13.3(0-76.7) | 16.7(0-72.4) |
| **High-reward/Risk selections, Run 2, %** | | |
| Mean(SD) | 15.5(15.9) | 15.5(16.6) |
| Median(range) | 10(0-53.3) | 10(0-60) |
| **High-reward/Risk selections, Run 3, %** | | |
| Mean(SD) | 10.8(15.0) | 12.9(16.3) |
| Median(range) | 3.6(0-53.3) | 6.7(0-60.7) |

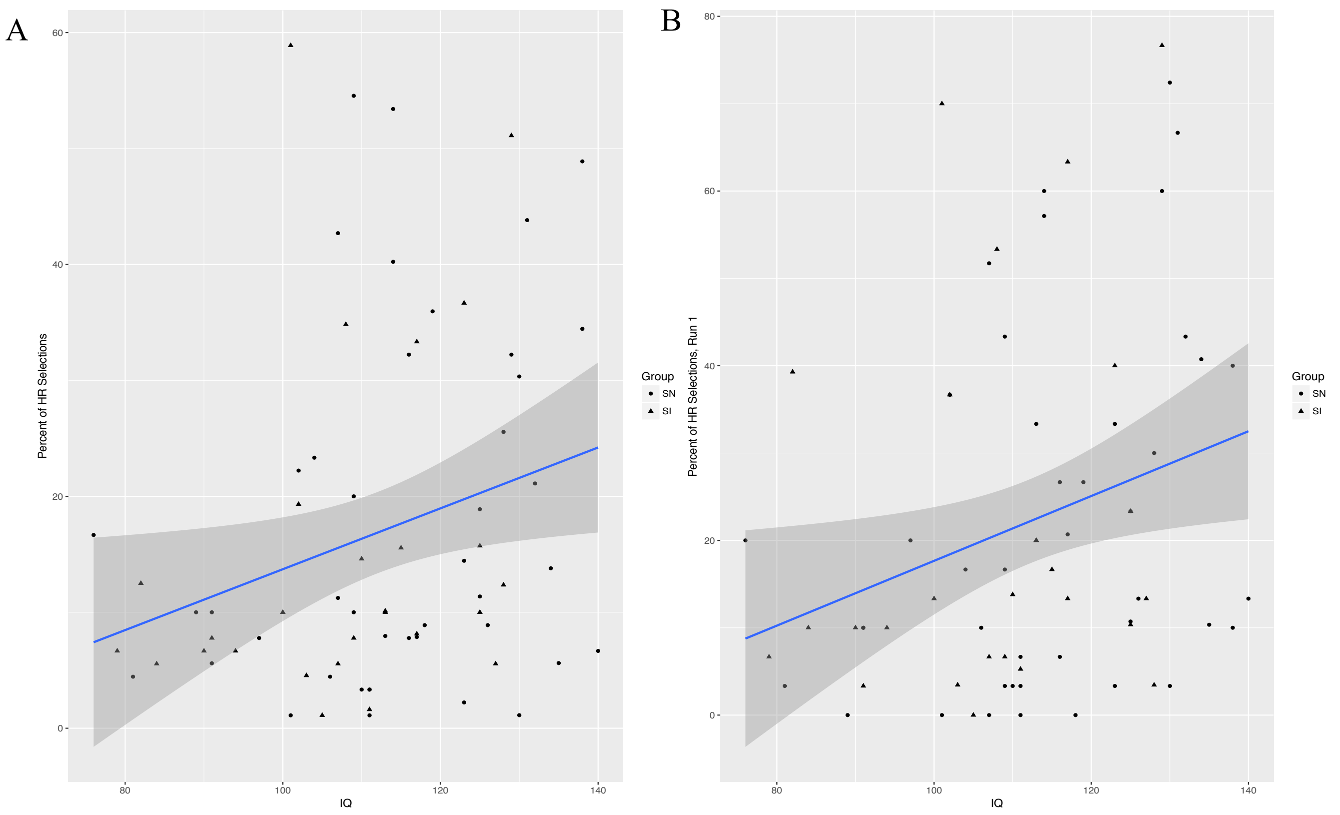

*Figure S1. Scatterplots showing the association between FSIQ in total group (N=70) with the percent of high-reward/risk (HR) decisions A) for all three runs of the WOF task and B) for the first run of the WOF task.*

*Table S3. Number of participants with zero high-reward/risk selections by run*

| | SI<br>*n*=27 | SN<br>*n*=43 |
|---|---|---|
| **No High-reward/risk selections, Run 1** | | |
| n(%) | 1(3.7) | 5(11.6) |
| **No High-reward/risk selections, Run 2** | | |
| n(%) | 3(11.1) | 6(13.9) |
| **No High-reward/risk selections, Run 3** | | |
| n(%) | 8(29.7) | 6(13.9) |

*Table S4. Mean response time (RT) for high-reward/risk selections by run*

| | SI<br>*n*=27 | SN<br>*n*=43 |
|---|---|---|
| **High-reward/risk selections RT (ms), Run 1** | *n*=26 | *n*=38 |
| Mean(SD) | 1.31(.45) | 1.26(.43) |
| Median(range) | 1.27(.55-2.39) | 1.29(.33-1.95) |
| **High-reward/risk selections RT (ms), Run 2** | *n*=24 | *n*=37 |
| Mean(SD) | 1.14(.31) | 1.3(.51) |
| Median(range) | 1.19(.61-1.7) | 1.33(.28-2.25) |

| **High-reward/risk selections RT (ms), Run 3** | *n*=19 | *n*=37 |
|---|---|---|
| Mean(SD) | 1.18(.37) | 1.22(.49) |
| Median(range) | 1.21(.59-1.98) | 1.17(.3-2.43) |

### S2. RESULTS

#### *S2.1 FMRI results: Within-group analyses*

In the selection phase (High-reward/risk > Low-reward/risk), the SN group demonstrated significant activation in the left insula. During the feedback phase (Win > Lose), SN adolescents demonstrated significant activation in the left putamen and superior frontal gyrus/middle frontal gyrus, and right precentral gyrus and cingulate gyrus (Table 4). Among adolescents reporting SU initiation (SI group), no results survived correction for multiple comparisons for either contrast of interest (High-reward/risk > Low-reward/risk or Win > Lose).

*Table S5. Results for SN group for selection (High-reward/risk > Low-reward/risk) and feedback (Win > Lose) phases of WOF task. Initial cluster defining threshold =*p*<0.001, k=10 voxels. Reported results survive FWE cluster-correction (*p*<.05).*

| **Region** | **BA** | **Cluster size** | **MNI coordinates** x | y | z | ***Z*** | ***t*** | **Corrected *p*-value (FWE)** |
|---|---|---|---|---|---|---|---|---|
| **High-reward/risk > Low-reward/risk** | | | | | | | | |
| L insular cortex | -- | 342 | -28 | 16 | -12 | 4.30 | 4.86 | .002 |
| **Win > Lose** | | | | | | | | |
| L putamen | -- | 1660 | -14 | 4 | -12 | 4.67 | 5.41 | .000 |
| R superior frontal gyrus | 9 | 508 | 20 | 38 | 48 | 4.54 | 5.21 | .000 |
| R precentral gyrus | 4 | 1091 | 2 | -22 | 72 | 4.53 | 5.20 | .000 |
| R cingulate gyrus | -- | 200 | 28 | 0 | 24 | 4.49 | 5.14 | .037 |
| L superior frontal gyrus/middle frontal gyrus | 8 | 671 | -24 | 34 | 50 | 4.48 | 5.12 | .000 |

*S2.2 FMRI Results: Between-group analyses without FSIQ as a covariate of no interest*

*Table S6. Summary of SN > SI cluster-level corrected results for Win > Lose contrast (without FSIQ as a covariate of no interest). Initial cluster defining threshold = p<0.001, k =10 voxels, FWE cluster-corrected at p < .05.*

| **Region** | **Cluster size** | **MNI coordinates** | | | ***Z*** | **Corrected *p*-value** |
|---|---|---|---|---|---|---|
| | | **x** | **y** | **z** | | |
| Right cingulate gyrus | 423 | 2 | 16 | 24 | 4.81 | .001 |

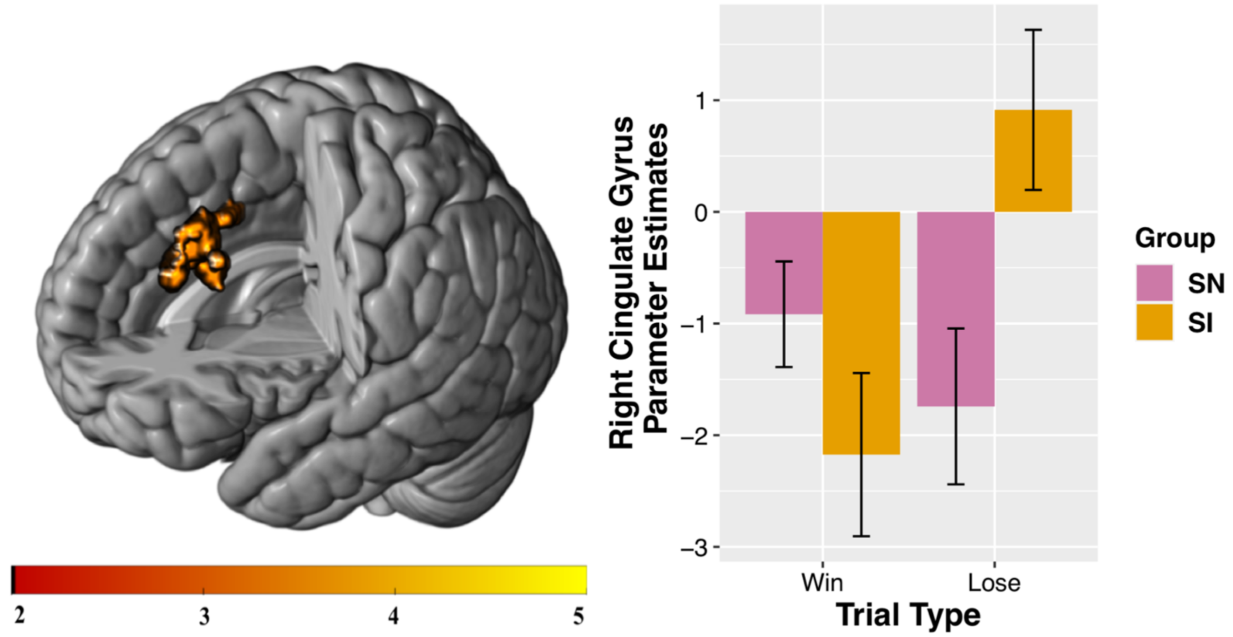

*Figure S2. Between-group results without FSIQ as a covariate of no interest. SN participants demonstrate greater activation relative to SI adolescents. Interaction charts depict mean parameter estimates and standard error for Win > Lose. Initial cluster defining threshold = p<0.001, k=10 voxels. Results survive FWE cluster-correction at* p<*.05.*

*S2.3 Characteristics of Wave 2 and Wave 3 SI participants*

*Table S7. Substance use reported by Wave 2 and Wave 3 SI participants*

| | **Wave 2 Initiators (~18-months follow-up)** (N = 12) | **Wave 3 Initiators (~36-months follow-up)** (N = 15) |
|---|---|---|
| **Mean age at initiation** | 14.79 (0.41) | 15.78 (0.70) |

| | | |
|---|---|---|
| **Substances reported** | Alcohol (11), marijuana (8), synthetic marijuana (4), spice (3), OTC (3), prescription painkillers (2), smoking tobacco (3), chewing tobacco (2) | Alcohol (13), marijuana (6), smoking tobacco (1), OTC (1) |
| **Polysubstance Users** | 9 | 4 |

*OTC = over-the-counter medications used to get high.*

*Table S8. Characteristics of Wave 2 and Wave 3 SI participants*

| | **SI (Wave 2)** *n* = 12 | **SI (Wave 3)** *n* = 15 | **Test statistic** | ***p*** |
|---|---|---|---|---|
| **Age at scan** | | | | |
| Mean(SD) | 12.98(.57) | 12.86(.65) | $t(25) = .50$ | .62 |
| **Sex** | | | $\chi^2(1) = 1.1$ | .30 |
| Females, n(%) | 8(67%) | 7(47%) | | |
| Males, n(%) | 4(33%) | 8(53%) | | |
| **PDS** | | | | |
| Mean(SD) | 2.7(.59) | 2.3(.64) | $t(25) = 1.7$ | .11 |
| Females | 2.9(.47) | 2.5(.80) | $t(13) = 1.2$ | .27 |
| Males | 2.2(.59) | 2.1(.41) | $t(10) = .52$ | .61 |
| **Race, n(%)** | | | $\chi^2(3) = 4.7$ | .19 |
| African American | 3(25%) | 3(20%) | | |
| Caucasian | 5(42% ) | 10(67%) | | |
| Hispanic/Latina/o | 1(8%) | 2(13%) | | |
| Other | 3(25%) | 0(0%) | | |
| **FSIQ** | | | | |
| Mean(SD) | 102.7 (14.2) | 111.7(13.6) | $t(25) = -1.7$ | .10 |
| **SES Index *z*-score** | | | | |
| Mean(SD) | -.25(1.15) | .20(.71) | $t(25) = -1.3$ | .22 |
| *Parental education, years, Mean (SD)* | 16.0(3.4) | 16.8(2.3) | | |
| *Household income, Median* | $50,000 - $74,999 | $1000,000 - $149,999 | | |
| **Family History** (FH+/-), n (%) *N* = 26 | N = 12 | N = 14 | $\chi^2(1)=.47$ | .50 |
| FHP | 4(33%) | 3(21%) | | |
| FHN | 8(67%) | 11(79%) | | |

*PDS = pubertal development scale, FSIQ = full-scale IQ, SES = socioeconomic, FHP = positive family history of alcohol/drug abuse, FHN = negative family history of alcohol/drug abuse.*

*S2.4 DUSI-R APD, DD task, and BIS/BAS: Wave 2 vs. Wave 3 SI participants*

Within the SI group, a one-tailed independent samples *t*-test of initial assessment/baseline DUSI APD revealed Wave 2 SI participants (mean: 23.15(12.59)) showed significantly higher scores compared to Wave 3 SI adolescents (mean: 15.44(9.55)) ($t(24) = 1.78$, $p = .04$)). Wave 2 and Wave 3 SI adolescents did not differ for DD, or for BIS/BAS subscales.

*Table S9. DUSI-R, DD, and BIS/BAS subscales for Wave 2 and Wave 3 SI participants*

| | **All SI**<br>**N = 27** | **SI**<br>**(Wave 2)**<br>**N = 12** | **SI**<br>**(Wave 3)**<br>**N = 15** | **Test statistic** | ***p*** |
|---|---|---|---|---|---|
| **DUSI-R APD** | | N = 12 | N = 14 | | |
| Mean(SD) | | | | | |
| *N = 26* | 19.0(11.5) | 23.1(12.6) | 15.4(9.5) | *t*(24) =1.78 | .044* |
| **DD, AUClogd** | N = 25 | N =11 | N = 14 | | |
| Mean(SD) | .54(.19) | .56(.21) | .52(.18) | *t*(23) = .44 | .66 |
| **BAS Drive** | 10.7(2.4) | 9.33 | 9.2(2.9) | *t*(25) =.14 | .89 |
| Mean(SD) | 11(6-15) | 9.5(7-12) | 8(6-15) | | |
| **BAS Fun-seeking** | 12.15(2.2) | 7.1(1.4) | 8.5(2.6) | *U* = 60.5 | .14 |
| Median(Range) | 13(5–14) | 7(5-9) | 7(5-14) | | |
| **BAS Reward Responsivity** | 17.78(1.6) | 7.1(1.4) | 7.3(1.8) | *t*(25) = -.40 | .69 |
| Median(Range) | 18(5-11) | 7(5-9) | 7(5-11) | | |
| **BIS** | 15 (3.11) | 15.7(3.2) | 15.1(3.1) | *t*(25) =.49 | .63 |
| Median(Range) | 15.3(11-23) | 15(11-23) | 15(11-22) | | |

*Note. Group comparisons for the BAS fun seeking scale used a Mann-Whitney U-test because assumption of normality for parametric testing was not met. DUSI-R APD used a one-tailed independent samples* t*-test. All other tests were two-tailed. * = p < .05. DUSI-R APD = Drug Use Screening Inventory, Revised, Absolute Problem Density; DD, AUC = delay discounting, area under the curve; BAS = Behavioral Activation System; BIS = Behavioral Inhibition System.*

*S2.5 WOF task behavior: Wave 2 and Wave 3 SI participants*

Initiation groups did not differ for percent of high-reward/risk selections ($Z$=70, $p$=.33). To determine whether Wave 2 compared Wave 3 participants demonstrated differences in response time (RT) for high-reward/risk versus to low-reward/risk selections, a two-way repeated measures ANOVA was used to examine the effect of group (Wave 2 SU Initiator vs. Wave 3 SU Initiator) and selection type (high-reward/risk vs. low-reward/risk) on RT. A main effect of selection type was found, with both groups demonstrating significantly slower RT in making high-risk compared to low-reward/risk selections ($F(1,25)=31.4$, $p<.000$). There was neither an effect of group ($F(1,25)=.39$, $p=.54$), nor a significant group × selection type interaction ($F(1,25)=.06$, $p=.81$).

*Table S10. WOF task behavior for Wave 2 and Wave 3 SI participants: descriptive statistics*

| | **All Initiators** | **Wave 2 Initiator** (~18-months follow-up) (*n*=12) | **Wave 3 Initiators** (~36-months follow-up) (*n*=15) |
|---|---|---|---|
| **High-risk/reward selections, %** | | | |
| Mean(SD) | 15(15) | 10.8(9.3) | 19.9(17.5) |
| Median (range) | 10(1-59) | 7.96(1.1-36.7) | 10.11(4.5-58.9) |
| **High-risk/reward selections RT (ms)** | | | |
| Mean(SD) | 1240(390) | 1200(410) | 1270(390) |
| **Low-risk/reward selections RT (ms)** | | | |
| Mean(SD) | 990(270) | 940(260) | 1030(280) |
| **All selections RT (ms)** | | | |
| Mean(SD) | 1110(320) | 1070(310) | 1150(320) |
